# Incorporating genome-based phylogeny and functional similarity into diversity assessments helps to resolve a global collection of human gut metagenomes

**DOI:** 10.1101/2020.07.16.207845

**Authors:** Nicholas D. Youngblut, Jacobo de la Cuesta-Zuluaga, Ruth E. Ley

**Author notes:** Corresponding author: Nicholas Youngblut.

## Abstract

Tree-based diversity measures incorporate phylogenetic or functional relatedness into comparisons of microbial communities. This can improve the identification of explanatory factors compared to tree-agnostic diversity measures. However, applying tree-based diversity measures to metagenome data is more challenging than for single-locus sequencing (*e*.*g*., 16S rRNA gene). The Genome Taxonomy Database (GTDB) provides a genome-based reference database that can be used for species-level metagenome profiling, and a multi-locus phylogeny of all genomes that can be employed for diversity calculations. This approach also allows for functional diversity measures based on genomic content or traits inferred from it. Still, it is unclear how metagenome-based assessments of microbiome diversity benefit from incorporating phylogeny or function into measures of diversity. We assessed this by measuring phylogeny-based, function-based, and tree-agnostic diversity measures from a large, global collection of human gut metagenomes composed of 33 studies and 3348 samples. We found tree-based measures to explain phenotypic variation (*e*.*g*., westernization, disease status, and gender) better or on par with tree-agnostic measures. Ecophylogenetic and functional diversity measures provided unique insight into how microbiome diversity was partitioned by phenotype. Tree-based measures greatly improved machine learning model performance for predicting westernization, disease status, and gender, relative to models trained solely on tree-agnostic measures. Notably, ecophylogenetic and functional diversity measures were generally the most important features for predictive performance. Our findings illustrate the usefulness of tree- and function-based measures for metagenomic assessments of microbial diversity – a fundamental component of microbiome science.

**Importance:** Estimations of microbiome diversity are fundamental to understanding spatiotemporal changes of microbial communities and identifying which factors mediate such changes. Tree-based measures of diversity, which consider species relatedness, are widespread for amplicon-based microbiome studies due to their utility relative to tree-agnostic measures. However, tree-based measures are seldomly applied to shotgun metagenomics data. We evaluated the utility of phylogeny, functional relatedness, and tree-agnostic diversity measures on a large scale human gut metagenome dataset to help guide researchers with the complex task of evaluating microbiome diversity via metagenomics.

## Introduction

Sequencing-based assessments of microbiome diversity are fundamental to the field of microbiome science. Metagenomic and 16S rRNA gene sequence-based estimations of human gut microbiome diversity have shown substantial differences among individuals due to disease state, diet, exercise, hygiene, and antibiotic usage (1). Measures of diversity differ in what data is utilized. For example, some only use taxon presence, while others incorporate abundances (2). Certain diversity measures such as Faith’s Phylogenetic Diversity (Faith’s PD) and UniFrac incorporate relatedness of taxa (3, 4), often via a tree inferred from ≥1 conserved phylogenetic marker, such as the 16S rRNA gene. These methods thereby leverage information ignored by tree-agnostic approaches, which assume that all taxa are equally related (*i*.*e*., star phylogeny measures) (2).

The importance of taxon relatedness in the assessment of diversity depends on the degree to which community assembly matches the pattern of microbial evolutionary/functional relatedness (5, 6). For example, if evolutionary relatedness corresponds strongly with phenotypic relatedness, specifically regarding phenotypes under selective pressure along the focal environmental gradient, then evolutionary relatedness is an informative prior for modeling the distribution of diversity across the gradient. Yet, phylogeny-based assessments of diversity may not always be more informative than tree-agnostic measures. For instance, phenotypes under selection may not be evolutionarily conserved due to recombination or convergent evolution. Studies of microbial phenotype evolutionary conservation suggest that this varies depending on the taxonomic clade and phenotype (7). Phylogeny-based diversity measures are also dependent on the accuracy of the phylogenetic inference, which may be questionable in the most common scenario for 16S rRNA-based surveys that usually rely on phylogenies inferred from only 1-2 hypervariable regions of the 16S rRNA gene (8). Even with these caveats, many studies have utilized phylogenetic measures of diversity to differentiate microbiomes by host disease state or other phenotype (9, 10). Moreover, phylogenies are successfully utilized in many recently published tools for microbiome data analysis, which include pseudo-count imputation (11), dataset augmentation (12), constrained ordination analysis (13), transforming compositional abundance data (14), and machine learning models for predicting host phenotypes (15, 16).

As the cost of sequencing has declined, shotgun metagenomics has risen in popularity relative to single-locus sequencing, especially since metagenomics provides a great wealth of information, including i) accurate species-level taxonomic classification and abundance estimation, ii) information on gene and metabolic pathway content, and iii) the ability to assemble genes and metagenome-assembled genomes (MAGs) (17–19). Recent work has shown that shallow sequencing depths can provide similar or greater coverage of microbial diversity compared to 16S rRNA sequencing (20). However, generating a phylogeny from shotgun metagenome data is inherently challenging, since sequences originate from all genomic locations instead of a single locus (21). Various methods exist for extracting 16S rRNA gene sequences, other single loci, or multi-locus data from metagenome reads (*e*.*g*., EMIRGE, MATAM, AMPHORA2, PhyloSIFT, and MetaPhlAn2), but using only single- or multi-locus information excludes much of the data, limiting the detection sensitivity for less common taxa (22–26). Alternatively, assembling MAGs enables the construction of multi-locus phylogenies from all assembled genomes, but very high sequencing depths are required to assemble rarer taxa in diverse microbial communities such as in soil or the human gut. Another approach is to map all reads to a database of genome assemblies (*e*.*g*., GenBank or RefSeq), which increases the amount of reads classified relative to single- or multi-locus approaches, but such databases usually lack careful curation of genome assembly quality, a standardized taxonomy, and multi-locus phylogenies for all reference genomes (27).

We recently created a pipeline for generating custom metagenome profiling databases from the Genome Taxonomy Database (GTDB) (27–29), which is a comprehensive database of *Bacteria* and *Archaea* genomes, which not only provides a coherent microbial taxonomy based on genome relatedness, but also includes multi-locus genome phylogenies for the reference species. Therefore, one can map all reads to the GTDB reference genomes to infer species-level abundances (*e*.*g*., with Kraken2) and then utilize a genome phylogeny of reference species for calculating alpha and beta diversity (30). Importantly, the multi-locus genome phylogeny will almost definitely be more robust and better-resolved than a phylogeny inferred from small, hypervariable regions of the 16S rRNA gene, or even the full-length gene sequence (31).

Using species-level reference genomes also enables “functional” based assessments of diversity (32), in which information on putative phenotypes is derived from genomic content of each reference (*e*.*g*., gene Pfam annotations). Moreover, high-level phenotypes (*e*.*g*., cell morphology, anaerobiosis, and spore formation) can be accurately inferred from low-level genomic content via machine learning methods (33), which allows for another, possibly more interpretable functional diversity assessment. Functional relatedness, whether derived from genomic content or predicted phenotypes can be integrated into existing tree-based diversity measures by representing the data as a dendrogram of functional relatedness. Such an approach has been scarcely used for 16S rRNA-based surveys, given the lack of direct phenotypic information obtainable from 16S rRNA sequences. Nevertheless, trait-based assessments of microbiome diversity that focused on a few key phenotypes have been employed with great effect in some circumstances (34, 35).

While promising, the approach of species-level metagenome profiling, followed by phylogeny- or function-based diversity calculation has not been robustly assessed and compared to tree-agnostic approaches that are often used for shotgun metagenome studies (*e*.*g*., Jaccard and Bray-Curtis distance measures). We therefore applied this methodology to a large, global human gut metagenome collection comprising 33 datasets and 3348 samples. We found that, in comparison to tree-agnostic measures, both phylogeny- and function-based measures of alpha and beta diversity improved our ability to discriminate metagenome samples based on westernization, disease status (*i*.*e*., healthy vs diseased), gender, and age. Moreover, ecophylogenetic and functional diversity measures, which are often not applied to metagenomic data, further resolved how these phenotypes partitioned gut microbiome diversity.

## Methods

### Data Retrieval

We retrieved publicly available human gut metagenomes from the Sequence Read Archive (SRA) between December 2019 and February 2020 (Table S1). Sample metadata was obtained from the curatedMetagenomicData v.1.17.0 Bioconductor package (36) and included according to the following criteria: i) shotgun metagenomes sequenced using the Illumina HiSeq platform with a median read length >95 bp; ii) with available SRA accession; iii) labeled as adults or seniors, or with a reported age ≥18 years; iv) without report of antibiotic consumption (*i*.*e*., no or NA); v) without report of pregnancy (*i*.*e*., no or NA); vi) non-lactating women (*i*.*e*., no or NA); vii) without report of gangrene, pneumonia, cellulitis, adenoma, colorectal cancer, arthritis, Behcet’s disease, cirrhosis or inflammatory bowel disease. Only forward reads were downloaded and further processed. The final dataset was composed of 3348 samples from 33 studies.

### Sequence processing and taxonomic profiling

We used the bbtools “bbduk” command and Skewer v0.2.2 (37) to trim adapters and quality-filter raw sequences. The “bbmap” command from bbtools was used to remove human reads mapping to the human genome hg19 assembly. We created quality reports for each step using fastqc v0.11.7 (https://github.com/s-andrews/FastQC) and multiQC v.1.5a (38). Filtered reads were subsampled to 1 million reads per sample and used to obtain taxonomic profiles using Kraken2 (30) and Bracken v2.2 (19). Custom databases of Bacteria and Archaea were created using Struo v0.1.6 (28) and based on the Genome Taxonomy Database (GTDB), Release 89.0 (“GTDB-r89”; available at http://ftp.tue.mpg.de/ebio/projects/struo/) (27).

### Genome phylogeny

The GTDB-r89 “Arc122” and “Bac120” multi-locus phylogenies were obtained from the GTDB ftp server (https://data.ace.uq.edu.au/public/gtdb/data/releases/release89/). The ape R package was used to merge the trees and prune them to the 23,360 species in the Struo-generated GTDB-r89 metagenome profiling database (28).

### Genomic content

Genes from from all selected GTDB-r89 reference genomes were annotated against UniRef90 (release 2019-10) via the *mmseqs search* function (-e 1e-3 --max-accept 1 --max-seqs 100 --num-iterations 2 --start-sens 1 --sens-steps 3 -s 6) from MMseqs2 (39). UniRef90 annotations were mapped to Clusters of Orthologous Groups (COG) and Pfam annotations via mapping files available from the HUMAnN3 database (17). We used the vegan R package (40) to apply the Bray-Curtis dissimilarity metric to the annotations-per-genome matrix to create a distance matrix of genome compositional relatedness. This distance matrix was clustered via the UPGMA algorithm to create a dendrogram, which was used for tree-based alpha and beta diversity metrics.

### Trait inference

We generated a Python v3 implementation of traitar (33) and used it to predict traits for all genomes based on pfam annotations (https://github.com/nick-youngblut/traitar3), with majority-rules (phypat+PGL model) used for classifying trait presence/absence. A dendrogram of trait relatedness was created as done with the COG/Pfam trees, except the Jaccard metric was used on per-genome trait presence/absence.

### Congruence of the genome phylogeny and trait similarity

Global congruence of the genome phylogeny, trait, and COG/Pfam similarity dendrogram was assessed via phytools::cospeciation with 100 permutations for the null model. Local congruence (*i*.*e*., per-clade) was assessed via Procrustes superimposition (vegan::procrustes) comparing the genome phylogeny patristic distance matrix versus the Jaccard distance matrix used to generate the trait similarity dendrogram. Due to memory limitation issues with the standard approach for converting a phylogeny to a patristic distance matrix in R (*i*.*e*., the “cophenetic” function in the ape R package can only process trees with fewer than ∼13,000 tips), we instead ran the procrustes analysis on 100 randomly pruned subtrees of 5000 tips each and used the mean residuals across all permutations for each taxon.

### Alpha diversity

All tree-agnostic measures were calculated with the vegan R package (vegan::diversity), Faith’s PD, mean pairwise distance (MPD), and mean nearest taxon distance (MNTD) were calculated with the PhyloMeasures R package (pd.query, mpd.query, and mntd.query functions, respectively) (41). Functional richness and evenness were calculated from all traits (*n* = 67) or COG categories (*n* = 23) per reference genome via the FD R package (FD::dbFD) (42). Table S2 lists and describes all diversity measures used for machine learning.

The lme4 and lmerTest R packages were used to fit linear mixed effects models with dataset as random effect and other variables as fixed effects; *F*-tests and *P*-values were determined via the Satterthwaite’s method (ANOVA Type II sum of squares). Cohen’s D was calculated with the effsize R package (effsize::cohen.d) (43). We adjusted *P*-values for multiple comparisons using the Benjamini-Hochberg method.

### Machine learning classifiers of host variables

Machine learning was performed with the mlr R package (44). We used random forests based on conditional inference trees, as implemented in the party R package (45). All hyperparameters were set to default, except ntrees = 500. Performance was evaluated via 5-fold cross validation with the Area Under the Curve (AUC) and F1 metrics. Train-test splits were blocked by study. Subsampling was used to balance the target variable, with the following undersampling ratios: westernization = 0.2, disease status = 0.33, gender = 0.66.

### Beta diversity analyses

Tree-agnostic weighted and unweighted intersample distances (Bray-Curtis dissimilarity and Jaccard index) were calculated using the vegan R package (vegan::vegdist), while tree-based metrics (weighted and unweighted UniFrac) were calculated with rbiom::unifrac. Principal coordinates analysis (PCoA) was applied to each distance matrix via stats::cmdscale. We used the vegan::envfit function to assess correlations of species abundances to each PCoA. We assessed PCoA ordination similarity via Procrustes superimposition (999 permutations).

### General data analysis

General data processing was performed with the tidyverse package in R (46). The R ggplot2 package was used for generating all plots (47). For all boxplots, box centerlines, edges, whiskers, and points signify the median, interquartile range (IQR), 1.5 × IQR, and >1.5 × IQR, respectively. All code used for this work is available on GitHub at https://github.com/leylabmpi/global_metagenome_diversity.

### Data availability

The genome phylogeny, trait table, COG/pfam gene annotation tables, and trait/COG/Pfam dendrograms for all species-genome representatives, are available at http://ftp.tue.mpg.de/ebio/projects/struo/GTDB_release89/.

## Results

### Dataset summary

Our combined human gut metagenome dataset consisted of 33 studies and a total of 3348 samples from 3011 individuals after filtering by required metadata fields and an adequate number of reads following quality control (101 ± 163 s.d. samples per study; Figure 1 & S1). The percent of metagenome reads classified to our custom GTDB-r89 Kraken2 database was high (mean of 80%), and tended to be lower for non-westernized populations.

**Figure 1.**
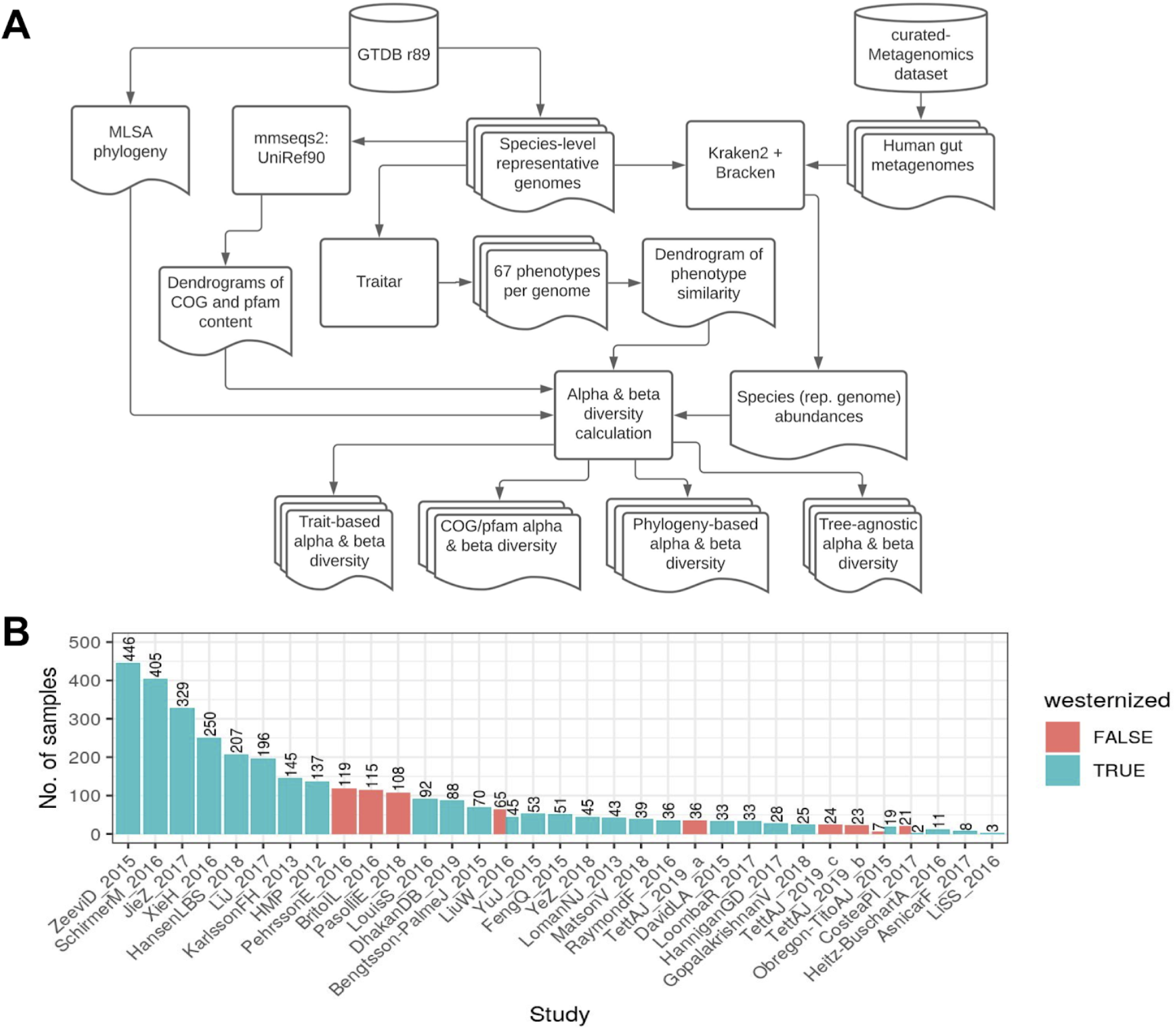
Methodological workflow and metagenome dataset summary. A) An overview of the workflow used. Cylinders denote public sequence databases (“r89” is “Release 89”), rounded squares denote analyses (*e*.*g*., running Kraken2), and squares with wavy bottoms denote output data files. B) The number of samples per dataset (a total of 3348 metagenomes) colored by westernization status. The text above each bar denotes the number of samples.

### Broad-scale incongruences between trait, gene annotation, and phylogenetic similarity

By using a genome reference dataset for taxonomic profiling, we could harness an existing genome phylogeny inferred from all reference genomes. To contrast phylogenetic and phenotypic relatedness, we inferred 3 more trees based on i) 67 traits inferred from all reference genomes via a machine learning method (Figure 2A), ii) the Pfam content of all reference genomes, and iii) the COG content of all genomes. Prior to utilizing the 4 trees for tree-based alpha and beta diversity assessments, we compared the congruence among trees to assess whether each could reveal different patterns of alpha and beta diversity. Pairwise congruence between the trees was measured via Procrustes superimposition, in which larger incongruences produce larger Procrustes residuals. Notably, residuals are on the per-taxon level, so we could assess how taxa in each clade (*e*.*g*., phylum) differed in phylogenetic and functional relatedness.

**Figure 2.**
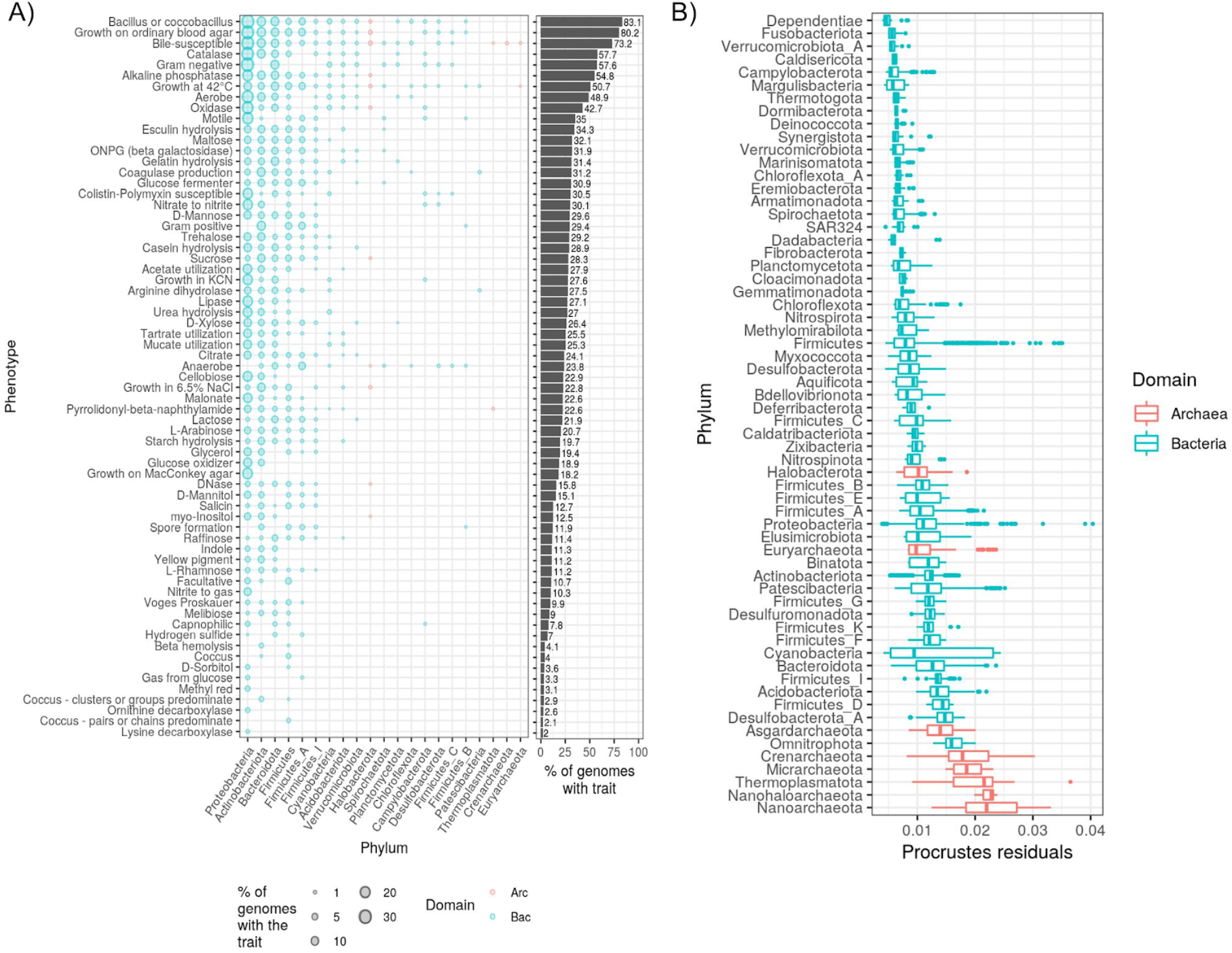
Similarity between phylogenetic and trait-based relatedness differs substantially among phyla. A) Traits inferred from each genome representative of each species, shown as the percent of all genomes in the phylum (left) or the total for all phyla (right). The numbers next to each column in the right panel denote the x-axis values. B) The boxplots show Procrustes residuals for each genome, grouped by phylum. Higher Procrustes residuals indicate more incongruence between phylogenetic and trait-based relatedness. For clarity, only phyla with ≥10 genomes are shown.

When comparing the genome phylogeny to the 3 functional trees, we observed similar incongruences, regardless of the functional tree (Figure S2). Given that the traits were inferred from Pfam content, the high correspondence between functional trees suggests that the high-level traits well-capture low-level variability in Pfam and also COG genomic content. Still, a few phyla did notably differ between the trait and gene annotation trees (*e*.*g*., *Elusimicrobia* and *Binatota*), possibly due to limited experimental phenotypic characterization and thus poor prediction accuracy of the target traits. For subsequent assessments of tree congruence, we focus on phylogeny versus traits, given the high congruence among the functional trees, and the ease of interpreting a limited number of high-level traits versus thousands of Pfam or COG features.

We found that the congruence between trait and phylogenetic similarity differed greatly across phyla (Figure 2B). The bacterial phyla *Dependentiae, Fusobacteriota*, and *Verrucomicrobiota A* were the most congruent between trait and phylogenetic similarity, while many of the archaeal phyla, including the *Crenarcheota, Thermoplasmatota*, and *Nanoarchaeota* were the most incongruent. The limited number of predicted traits for many of the archaeal clades may be a primary cause for this high incongruence (Figure 2A); however, the Pfam and COG trees show similarly high levels of incongruence (Figure S2), suggesting the non-exclusive possibilities that i) genomic content is not well characterized at the level of gene annotations and ii) rapid evolution of the variable component of the pan-genome relative to the core. Indeed, *Crenarcheota* were recently shown to be especially variable in phenotypes, as defined by overlap in clusters of orthologous groups (COG) functional categories (48).

*Firmicutes* and *Proteobacteria* showed the greatest variance in congruence, with many highly incongruent outlier species in both phyla. We note that variance was only weakly correlated with the number of species per phylum (Spearman’s ρ = 0.36), so the large variance among *Firmicutes* and *Proteobacteria* was not simply due to the clade size. An inspection at the family level revealed that the *Firmicutes* outliers belonged to *Enterobacteriaceae*, while the *Mycoplasmoidaceae* and *Metamycoplasmataceae* families were the largest outliers in *Proteobacteria* (Figure S3). *Euryarchaeota* trait-phylogeny congruence was relatively high for an archaeal phylum; however, the *Methanosphaera* genus comprised many highly incongruent outliers. Overall, our findings show that trait and phylogenetic similarities are only partially congruent and would thus likely describe different aspects of microbiome diversity when applied to tree-based diversity measures.

### Tree-based alpha diversity improves population differentiation

We calculated alpha diversity for all 3348 metagenome samples with 3 measures: the number of observed taxa, the Shannon Index, and Faith’s Phylogenetic Diversity (Faith’s PD) calculated with the genome phylogeny and each functional tree. Metagenomes were subsampled to 1 million reads prior to metagenome profiling; thus, alpha diversity estimates should not be biased by sampling depth. All tree-based alpha diversity measures clearly separated metagenome samples based on westernization status, while such a separation was less discernible when using the Shannon Index or the number of observed species (Figure 3A & 3B). When assessing samples with westernization status, age, and gender metadata (*n* = 1843), we also found that the tree-based measures more clearly differentiate groups along each variable (Figure S4). Indeed, linear mixed effects models produced substantially higher *F* values and more significant associations (adj. *P* < 0.01) for tree-based measures versus the Shannon index, in regards to westernization status, age, and gender (Figure 3C). In particular, *F* values were much higher for westernization when using Faith’s PD with the genome phylogeny. These results suggest that westernized and non-westernized individuals differ most strongly by the composition of distantly related taxa (*e*.*g*., different phyla), while high-level traits and lower-level genomic content is more similar among the human populations. Disease status was not significantly associated with any diversity measure, possibly due to disease-specific differences in the direction and magnitude of change in alpha diversity compared to healthy individuals.

**Figure 3.**
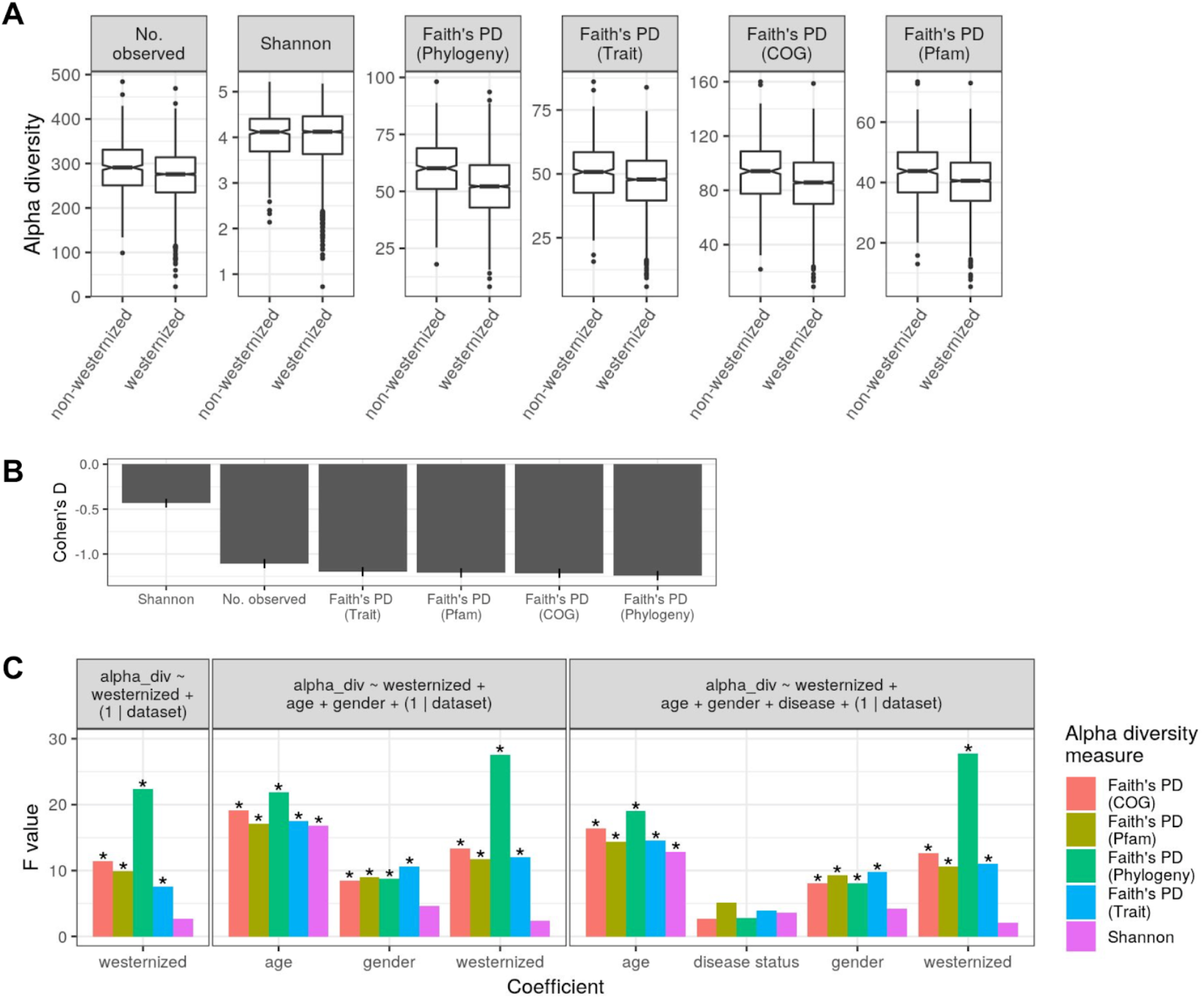
Tree-based alpha diversity better differentiates samples by key host phenotypes. A) Boxplots of alpha diversity metrics calculated for all samples (*n* = 3348) in all datasets (*n* = 33), grouped by westernization. “Phylogeny”, “Trait”, “COG”, and “Pfam” denote which tree was used to calculate Faith’s PD. B) Cohen’s D effect sizes for the difference in alpha diversity between non-westernized and westernized samples. The line ranges denote 95% confidence intervals. C) Linear mixed effects model results for assessing the association between alpha diversity and phenotypes while accounting for inter-dataset batch effects. Asterisks denote significant associations (adj. *P* < 0.01). Age was log_2_-transformed. Facet labels denote the model design (“disease” is disease status: healthy/diseased). The left facet is on all samples (*n* = 3348) in all datasets (*n* = 33). The middle facet is filtered to samples that have data on gender and age (number of samples = 1843; number of studies = 17). The right facet is filtered to samples that have data on gender, age, and disease status (number of samples = 1413; number of studies = 15).

To resolve how the choice of diversity measure influenced per-clade estimations of diversity, we applied our mixed effects model analysis on alpha diversity calculated for each individual taxonomic family (Figure S5). For all diversity measures, the *Bacteroidales* family F082 was most strongly associated with westernization, and the strength of association was very consistent among measures. The F082 family comprises 29 species, but only 4 were detected across the samples, and all belonged to one genus also named F082. In contrast, many of the other families associated with westernization differed in their strength among the diversity measures (Figure S5A). For instance, the association of *Treponemataceae* was substantially weaker for the Shannon Index versus the tree-based measures. This inconsistency among diversity measures was also observed for associations between family-level diversity and gender or age. Overall, *Akkermansiaceae* showed the strongest association with gender, although the association was substantially weaker for Faith’s PD based on the genome phylogeny (Figure S5), suggesting incongruences between functional and taxonomic diversity. Notably, *Methanobacteriaceae* alpha diversity was most strongly associated with age, along with *Butyricicoccaceae*, but the association strength was much lower when measuring diversity via phylogeny-based Faith’s PD versus the other diversity measures (Figure S5C). These examples show that fine taxonomic level diversity estimations can differ substantially depending on which aspects of diversity are emphasized: phylogenetic relatedness, trait relatedness, gene compositional similarity, or simply taxon abundances.

### Ecophylogenetic measures provide unique insight into community assembly

Phylogenies have been widely used in community ecology to investigate community assembly via such ecophylogenetic measures as the mean pairwise distance (MPD) and mean nearest taxon distance (MNTD) (49). These tests assess whether the distribution of taxon relatedness in each sample significantly differs from a permuted null model, which can provide insight about how selection pressures, competition, and other mechanisms affect community assembly. In general, comparing MPD to MNTD can provide insight into the relatedness of taxa in each community, given that MPD is influenced most by ancient diversification events, while MNTD is more sensitive to recent speciation events. We calculated MPD and MNTD for the phylogenetic, COG, Pfam, and trait trees. Similar to our previous alpha diversity analysis, linear mixed effects models showed many significant associations (adj. *P* < 0.05) with westernization, disease status, age, and gender (Figure 4A). The significance and strength of the associations differed substantially between the MPD and MNTD measures, and this was often dependent on the tree used. MPD was significantly associated with westernization, regardless of which tree was used, while significance was more variable for MNTD. Overall, gender and disease status were not significantly associated with MNTD, regardless of the tree used. These results indicate that westernization, age, gender, and disease status differ in how they partition gut microbiome diversity, both in regards to taxonomy versus phenotype and also the taxonomic/functional resolution in which partitioning occurs.

**Figure 4.**
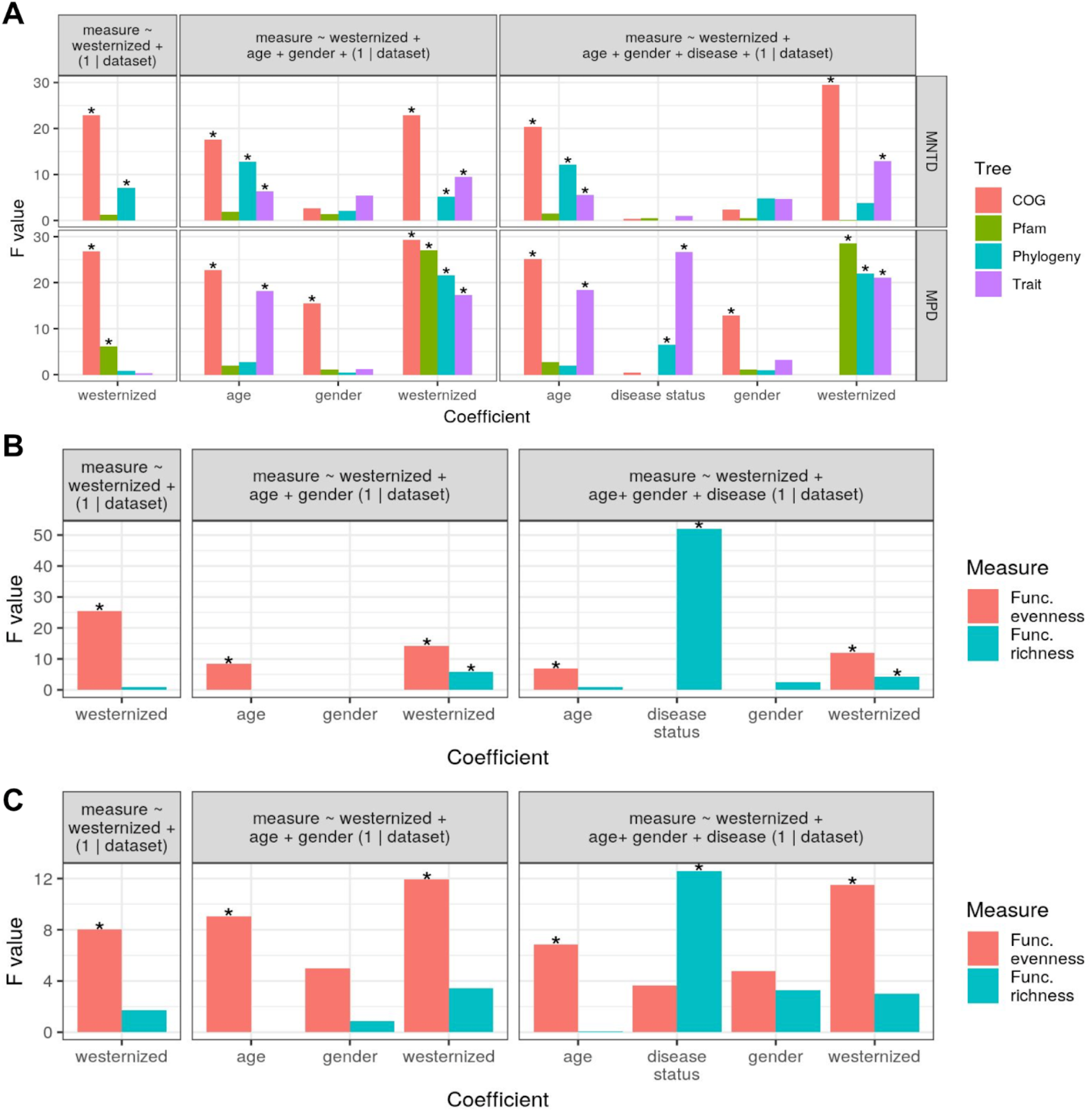
Ecophylogenetic and functional diversity (FD) measures resolve host phenotypes to varying degrees. Linear mixed effects model results for testing the association between host phenotypes and A) MNTD and MPD, B) FD based on 67 traits inferred from microbial genomic content, C) FD based on genomic content aggregated into 23 COG functional categories. Asterisks denote significant associations (adj. *P* < 0.05). Age was log2-transformed. Facet labels denote the model design (“disease” is disease status: healthy/diseased). The left facet is on all samples (*n* = 3348) in all datasets (*n* = 33). The middle facet is filtered to samples that have data on gender and age (number of samples = 1843; number of studies = 17). The right facet is filtered to samples that have data on gender, age, and disease status (number of samples = 1413; number of studies = 15).

### Functional evenness more strongly differentiates populations than functional richness

Our microbiome dataset consisted of per-taxon functional information in the form of predicted traits and COG/Pfam content of each genome. Therefore, we could investigate established community ecology measures of functional diversity (FD), which are not commonly used for microbiome data (32). Existing algorithms do not scale well to 1000’s of taxa with 100’s or 1000’s of phenotypes, so we limited our analysis to the 67 predicted traits and aggregating COGs to the 23 COG functional categories (*e*.*g*., “L” = Replication and repair). We calculated both functional evenness and functional richness; linear mixed effect models generally showed stronger and more numerous significant associations (adj. *P* < 0.05) for functional evenness, regardless of whether traits or COG categories were used (Figure 4B & 4C). We found that westernization, age, and gender were associated with functional evenness. In contrast, functional richness was only associated with disease status and westernization. Westernization was only associated with trait-based richness, but not richness of COG categories, while disease status was strongly associated with both calculations of functional richness. These findings suggest that functional evenness rather than diversity best explains age, gender, and westernization, while functional richness strongly explains disease status. Indeed, boxplots of FD values showed that functional richness was markedly higher among diseased individuals (Figure S6).

### Tree-based diversity measures improve random forest model performance

To assess the usefulness of various diversity measures for machine learning-based classification of host phenotypes, we trained binary random forest (RF) classification models to predict westernization, disease status, or gender. We trained models either with just tree-agnostic diversity measures (*n* = 4) or also including tree-based and FD measures (*n* = 32) to test whether tree-based measures increase the predictive capacity. We note that some diversity measures (*e*.*g*., Simpon’s Index and RaoQ) were not included in our previous mixed model analyses for the sake of clarity.

AUC and F1 performances across 5-fold cross validations were significantly higher when the tree-based diversity measures were included (Kruskal-Wallis, adj. *P* < 0.01), thus demonstrating the usefulness of tree-based measures for predicting phenotypes. The models trained on all measures showed performances comparable to similar microbiome-based machine learning models reported in the literature (50–52), with mean AUC values of 0.92, 0.82, and 0.67 for classifying westernization, disease status, or gender, respectively. We note that subsampling was used for each train-test evaluation in order to reduce overfitting resulting from dataset imbalances (see Methods).

Feature importance differed for each model trained on all diversity measures, but none of the tree-agnostic measures were in the top 10 for any model (Figure 5B-D). The general type of diversity measure (*e*.*g*., ecophylogenetic or FD) and data used for calculating diversity (*e*.*g*., phylogeny or traits) also differed in rank importance among models: all phylogeny-based measures (Faith’s PD, MNTD, and MPD) were in the top 10 for westernization or disease status classifiers, while only Faith’s PD was in the top 10 for the gender classifier. Notably, most of the top features were ecophylogenetic and FD measures, indicating that these seldomly-used measures may be broadly useful for microbiome-phenotype machine learning applications. Overall, the variability in rank-importance of diversity measures across models indicates that different measures help explain distinct aspects of community assembly in relation to particular host phenotypes, and thus incorporating multiple diversity measures may be a generalizable approach to phenotype classification tasks.

**Figure 5.**
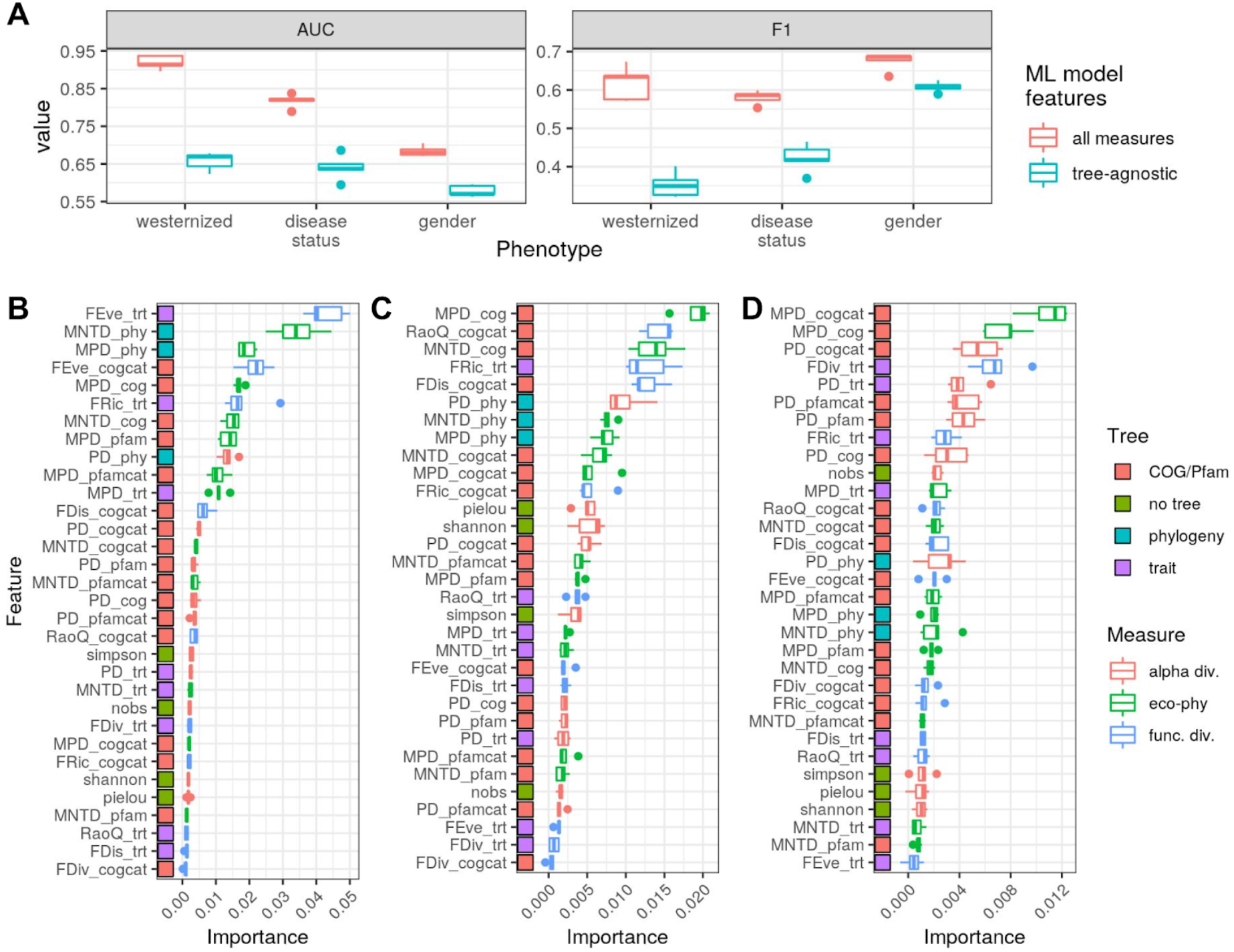
Tree-based diversity measures considerably improve random forest model prediction accuracy. A) Random forest (RF) model performances (AUC and F1) across all 5 cross validation (CV) folds for either all diversity measures (red) or just tree-agnostic measures (blue). The “all measures” performances were all significantly higher than the “tree-agnostic” performances (Kruskal-Wallis, adj. *P* < 0.01). The feature importances across CV folds for RF model predictions trained to predict B) westernization, C) disease status, or D) gender are shown via boxplots. The number of samples per model differed due to missing phenotype values: westernization = 3348, disease status = 2755, and gender = 2154. The colored boxes denote which tree was used for the diversity measure (“phylogeny” is the genome-based phylogeny), while the boxplot colors denote the type of diversity measure. “cogcat” and “pfamcat” are COG and Pfam annotations aggregated by general category (*n* = 23 and *n* = 609, respectively). See Methods for a description of each feature.

### More variance is explained by beta diversity measures incorporating phylogenetic or trait relatedness

We calculated beta diversity on all metagenome samples with 4 metrics: Bray-Curtis, Jaccard, and weighted/unweighted UniFrac. For both UniFrac measures, we used all 4 trees: the genome phylogeny and the 3 functional trees. Principal coordinate analysis (PCoA) revealed substantially differing amounts of variance explained by the top 2 PCs (Figure 6A). Indeed, pairwise comparisons of the ordianations via Procrustes superimposition showed that all ordinations significantly differed (adj. *P* < 0.05), but there was only minor differentiation between the tree-agnostic measures (Bray-Curtis and Jaccard) and also among the unweighted UniFrac measures (Figure S7). Phylogeny-based weighted UniFrac calculated differed most substantially from other measures, with an average effect size of 0.4. The results indicate that each measure can reveal unique aspects of how communities differ from one another, as is often currently done via comparing Bray-Curtis to Jaccard measures to determine the relative impact of abundant and rare taxa.

**Figure 6.**
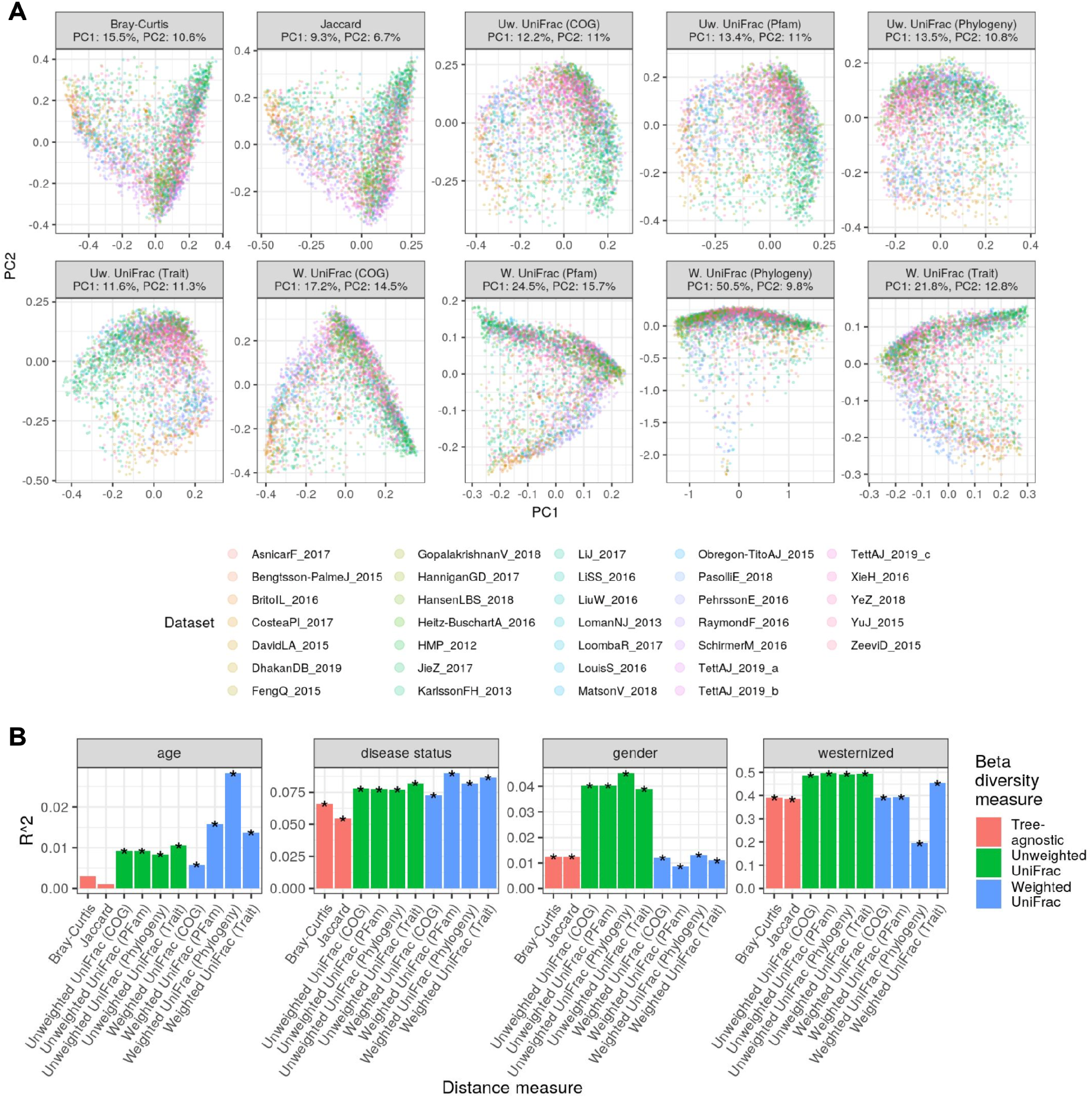
Beta diversity measures differ in their associations with host phenotypes. A) PCoA ordinations for each beta diversity measure. The facet labels state the percent variance explained by PC1 and PC2. B) EnvFit correlational analysis of age, gender, disease status (healthy/diseased), and westernization versus the top 2 PCoA PCs (*n* = 1438). Asterisks denote significant associations (adj. *P* < 0.05).

Given this variability among beta diversity measures, we tested whether host phenotypes were better resolved by certain measures. We correlated host age, gender, disease status, and westernization to the top 2 PCs of each ordination, which revealed differences in the significance and strength of association among the measures, depending on the phenotype (Figure 6B). All measures were significantly associated with all phenotypes (adj. *P* < 0.05), except for the tree-agnostic measures and age. In general, association strengths were higher for tree-based measures relative to tree-agnostic measures. The strongest associations were to westernization, but association strength (R^2^) ranged from 0.19 to 0.5, with weighted UniFrac measures generally the strongest, and phylogeny-based weighted UniFrac substantially lower than all others. All unweighted UniFrac measures also showed a substantially stronger association to gender than the other measures, but all measures were rather weakly associated (R^2^ range of 0.01-0.04). In contrast to westernization, phylogeny-based weighted UniFrac was most strongly associated with age, but as with gender, all associations were weak (R^2^ range of 0.01-0.03). Correlation strength was more consistent for disease status, with an R^2^ ranging from 0.05 to 0.09. In summary, these results indicate that tree-based beta diversity measures, as with tree-agnostic measures, can be useful for explaining phenotypic variation across human cohorts.

### Tree-based measures emphasize differences in certain taxa

Our results show that the beta diversity measures differ in which aspects of inter-sample taxonomic composition are emphasized (Figure S7). We therefore sought to determine which taxa contribute most to the beta diversity values for each measure. For the tree-based measures, we focused on the genome phylogeny and trait-based tree to simplify the analysis. We correlated taxon abundances with the top 3 PCs of each PCoA (Figure 7A), which revealed that the tree-based weighted UniFrac measures differentiated samples based on the abundances of species belonging to *Lachnospiraceae* (*Firmicutes A*), *Bacteroidaceae* (*Bacteroidota*), and *Enterobacteriaceae* (*Proteobacteria*). In contrast, the tree-agnostic measures most strongly discerned samples differing in species just within *Bacteroidaceae* (*Bacteroidota)*. Specifically, the top PCs for Bray-Curtis and Jaccard correlate strongly with the *Bacteroidaceae* genera: *Bacteroidetes, Bacteroidetes B*, and *Prevotella* (Figure S8). Unlike the weighted UniFrac measures, both unweighted UniFrac measures lacked a strong correlation with *Enterobacteriaceae*, but they did uniquely discern *Oscillospiraceae* (*Firmicutes A*) and *Ruminococcaceae* (*Firmicutes A*).

**Figure 7.**
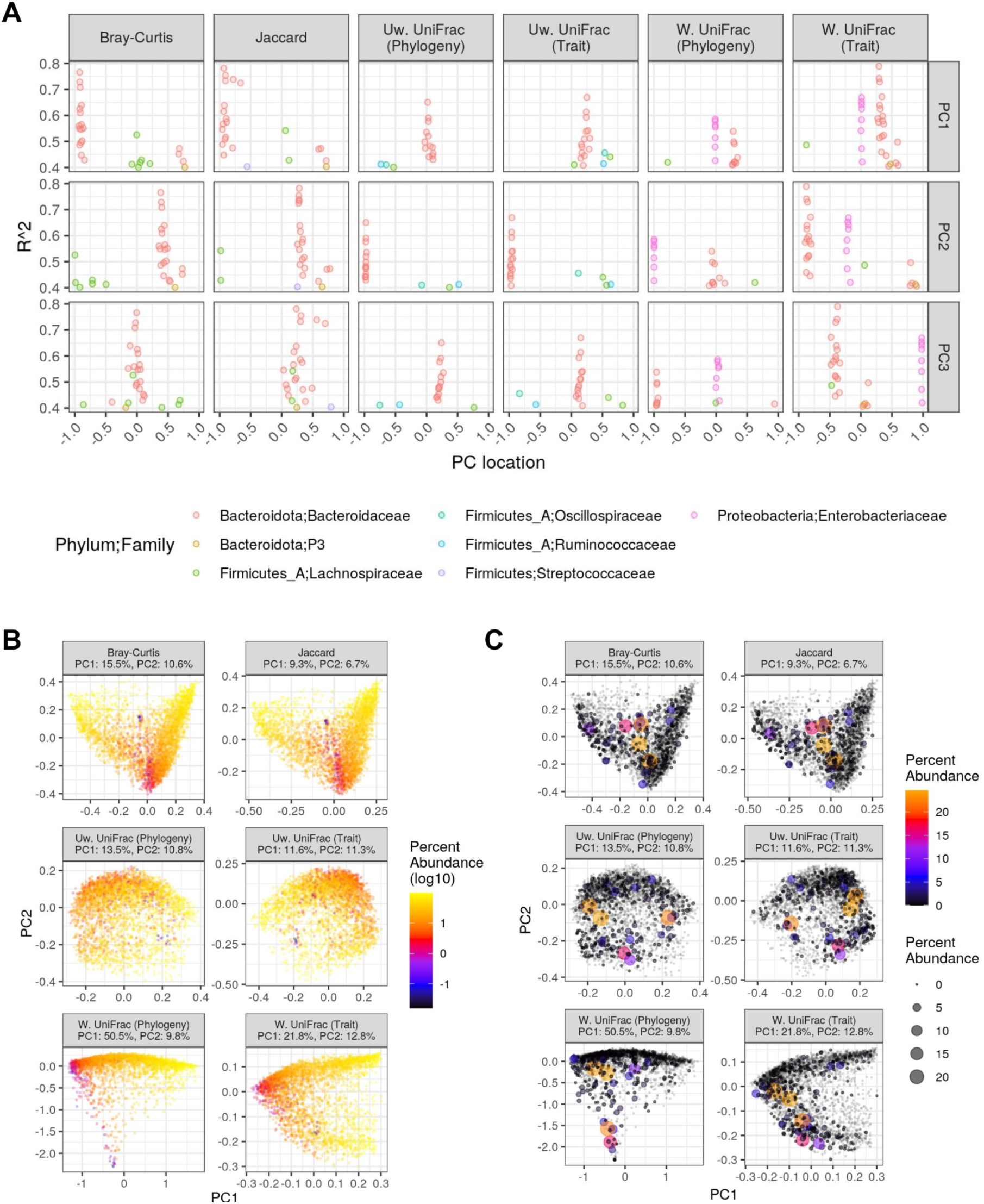
Phylogeny- and trait-based beta-diversity metrics emphasize inter-sample differences in certain taxa that are not emphasized by star-phylogeny metrics. A) Correlations between individual species (points) and the top 3 PCs in the PCoA ordinations shown in Figure 6. The x-axis denotes the direction of the correlation along the PC (*i*.*e*., where the taxon abundance is highest), and the y-axis denotes the effect size. For clarity, only species with the top 20 highest effect sizes across all beta diversity metrics are shown. The PCoA ordinations shown in B) and C) are the same as in Figure 6, but samples are colored by the abundance of the *Bacteroidaceae* family (*Bacteroidota* phylum) and *Enterobacteriaceae* (*Proteobacteria* phylum), respectively. Note that abundance is not log_10_-transformed in C), and point size also represents Enterobacteriaceae abundance in order to emphasize the few samples with relatively high *Enterobacteriaceae* abundances. Grey points indicate samples completely lacking *Enterobacteriaceae*.

To help illustrate these clade-level differences among the beta diversity measures, we mapped the abundances of these focal clades onto each PCoA ordination. As denoted by our correlation analysis, *Bacteroidaceae* was highly abundant at both ends of PC1 for Bray-Curtis and Jaccard, while its abundance was lowest at the center of the PC (Figure 7B). Conversely, *Bacteroidaceae* was only highly abundant on half of PC1 for both weighted UniFrac measures. In contrast to *Bacteroidaceae, Enterobacteriaceae* was only detectable in 350 samples, with only 28 samples having >1% abundance (Figure 7C). Trait-based weighted UniFrac best partitioned the samples with high versus low levels of *Enterobacteriaceae* (Figure 7A & 7C), while phylogeny-based weighted UniFrac also partitioned these samples well, especially along PC2. Plotting *Lachnospiraceae, Oscillospiraceae*, and *Ruminococcaceae* abundances on the PCoA ordinations confirmed the correlation analysis, in which *Lachnospiraceae* abundance correlates rather well with PC1 and PC2 of all ordinations, while the *Oscillospiraceae* and *Ruminococcaceae* abundances best correlate the the top PCs for both unweighted UniFrac measures (Figure S9).

We also correlated alpha diversity with the PCoA PCs but found substantially weaker associations (R^2^ < 0.21 for all measures). Still, gradients of diversity were somewhat apparent across the ordinations, regardless of the diversity measure (Figure S10).

## Discussion

Shotgun metagenomics will continue to increase in popularity as the cost of sequencing declines and methods for processing and interpreting metagenomic data continue to develop. A major challenge is to fully harness the heterogeneous sequence data generated by metagenomics, which is vastly more complex than 16S rRNA gene sequences or other single-locus datasets. Measuring community diversity from such heterogeneous data is not straight-forward, and it is often unclear what measures of diversity are most appropriate for metagenome data. Here, we have assessed a method of microbiome diversity measurement by using metrics that incorporate a multi-locus phylogeny, traits inferred from genomic content, and 2 categorizations of genomic content: Pfam and COG (Figure 1). This approach is possible due to using a reference database solely comprising genomes, rather than single locus reference databases such as SILVA or GreenGenes (53, 54), as is commonly used for 16S rRNA-based analyses (2). Our method is not computationally demanding for pre-characterized reference genomes, generalizable to a wide range of microbiome studies, and flexible in regards to which tree-based measures and to which genome characteristics are used (*e*.*g*., genes, inferred traits, or experimentally verified phenotypes).

Tree-agnostic measures of alpha and beta diversity do not require additional information beyond incidences or abundances of taxonomic or functional units, and the lack of direct comparisons between tree-agnostic and tree-based measures among existing metagenomic studies calls into question whether tree-based measures are necessary. Therefore, we directly compared the ability of tree-based and tree-agnostic diversity measures to explain phenotypic variation across a large metagenomic dataset, comprising 3348 samples and 33 studies. While the number of phenotypes was limited and the dataset solely comprised human gut metagenomes, the broad scope of the dataset suggests that our results will generalize to other biomes and phenotypes. Moreover, we sought to minimize the influence of technical factors such as study effects, sequencing depth, or unbalanced groups in our analyses. Our assessments of both alpha and beta diversity measures showed that tree-based measures generally explained phenotypes better or comparable with tree-agnostic approaches. For instance, all tree-based measures were significantly associated with westernization, gender, and age, while the tree-agnostic measure was only associated with age (Figures 3 & 6). Significance and strength of the associations varied by tree and phenotype, indicating incongruences between phenotypic and functional variation across populations. For example, westernization was better explained by phylogeny-based Faith’s PD compared to functional tree-based measures, indicating that the westernization status of a subject is better reflected in the taxonomic composition of their microbiome than in the metabolic potential encoded by the microbial community (*e*.*g*., Pfam annotations).

We illustrated how broad-level comparisons of diversity can provide insights into the components of microbiome assembly that differed by phenotype. First, we compared tree congruences in order to reveal which phyla and families are most congruent between evolutionary and functional relatedness (Figures 2, S2, & S3). While incongruences between the phylogeny and inferred traits could be a result of trait mis-classifications, similar discrepancies were observed between the phylogeny and COG/Pfam-based trees; thus, the high-level traits appear to capture well lower-level genomic content variation. Still, traits and genomic content differed to some extent, which is why we included all functional trees in subsequent analyses. Second, we performed mixed model analyses of alpha diversity for each taxonomic family to illustrate how diversity-phenotype associations can vary at the per-clade level due to incongruences between phylogenetic and functional relatedness (Figure S5). Third, when we assessed which taxa most strongly associated with the beta diversity patterns discerned from each measure (Figure 7), we found that tree-agnostic measures generally emphasized within-clade compositional differences among abundant and prevalent taxa, while tree-based measures emphasized inter-phylum differences among less abundance and lower prevalence clades. For instance, Bray-Curtis and Jaccard emphasized compositional differences within the *Bacteroidaceae*, which is a prevalent and abundant clade in the human gut. In contrast, both the phylogeny and trait-based measures accentuated differences between *Enterobacteriaceae and Bacteroidaceae*, which not only belong to different phyla, but also the former is much less prevalent than the latter. Of course, such correlational analyses tend to lack the sensitivity to assess compositional differences of rare taxa that are likely emphasized by the unweighted UniFrac measures, but this example of *Enterobacteriaceae* versus *Bacteroidaceae* highlights how tree-based and tree-agnostic diversity measures can reveal differing patterns of microbial diversity.

Beyond our direct comparison between tree-agnostic and tree-based diversity measures, we utilized diversity measures that can only be applied to tree-based data or per-taxon functional data (Figure 4). These analyses provided additional insight into how gut microbial diversity was partitioned across the human cohort. For example, comparison of MPD and MNTD measures, which emphasize different aspects of taxon relatedness in each community, showed that coarse phylogenetic and functional relatedness varied by phenotype, rather than taxa closely related by evolution or function (Figure 4). While MPD was generally more associated with each host phenotype relative to MNTD, some phenotype-diversity associations were only significant for MNTD. Age was significantly associated with phylogeny-based MNTD, while MPD was not, suggesting that age is linked to changes in closely related taxa. Our assessments of FD showed that westernization and age significantly differed in functional evenness, but less so for functional richness. In contrast, disease status was strongly associated only with functional richness, with a substantial increase of richness among diseased samples (Figure S6). While previous studies have shown disease-specific changes in taxonomic alpha diversity (55), FD has been less well-studied in this context, and usually function is assessed for all genes and/or metabolic pathways (*e*.*g*., via HUMAnN2) instead of per-taxon functional diversity, as performed in this study (17, 56). The overall increase of functional richness across all diseases may be the result of a consistent enrichment of certain high-level function categories in the microbiome, such as aerobiosis and antibiotic resistance.

By combining standard alpha diversity measures, such as Shannon index or Faith’s PD, with ecophylogenetic and FD measures, we were able to train phenotype-predicting machine learning models with accuracies comparable to the state-of-the-art (Figure 5). Notably, the most important features for predicting westernization, disease status, and age were ecophylogenetic and FD measures, rather than standard alpha diversity measures. Of the standard measures, tree-agnostic measures tended to have low predictive importance, and models solely comprising these measures performed significantly worse than models trained on all measures (Figure 5). Combining ecophylogenetic, FD, and standard diversity measures with taxon/gene/pathway abundances may produce models with even higher predictive performance, especially when combined with feature selection procedures to reduce the number of features relative to observations (50). More work is needed to evaluate whether the inclusion of many varying alpha diversity measures in microbiome-phenotype applications will generalize across different microbiome datasets. Still, the high model performances that we obtained when training on a broad dataset with large inter-study variation suggests a general applicability.

In summary, we have illustrated how tree-based and per-taxon functional diversity measures can be useful for metagenomic data analysis. Trees based on evolutionary or functional relatedness enable many additional assessments of diversity, especially given the variety of existing ecophylogenetic and FD measures. Inclusion of these additional measures may be fruitful for machine learning applications on metagenomic datasets. More generally, synthesizing phylogenetic and functional diversity may lead to the development of new, more encompassing ecological theory (57).

## Author contributions

Author contributions: N.D.Y. and J.dlC. designed the research; N.D.Y. and J.dlC. performed research; N.D.Y. analyzed data; and N.D.Y., J.dlC., and R.E.L. wrote the paper.

## Acknowledgements

We thank Albane Ruaud and Taichi Suzuki for providing important feedback on this work.

