## Supplemental Materials for "Incorporating genome-based phylogeny and functional similarity into diversity assessments helps to resolve a global collection of human gut metagenomes"

Nicholas D. Youngblut*[,1], Jacobo de la Cuesta-Zuluaga[1], Ruth E. Ley[1]
[1]Department of Microbiome Science, Max Planck Institute for Developmental Biology, Max Planck Ring 5, 72076 Tübingen, Germany

#### Supplemental Tables

**Table S1.** Metadata for all metagenome samples used in this work.

**Table S2.** All diversity measures used for machine learning.

### Supplemental Figures

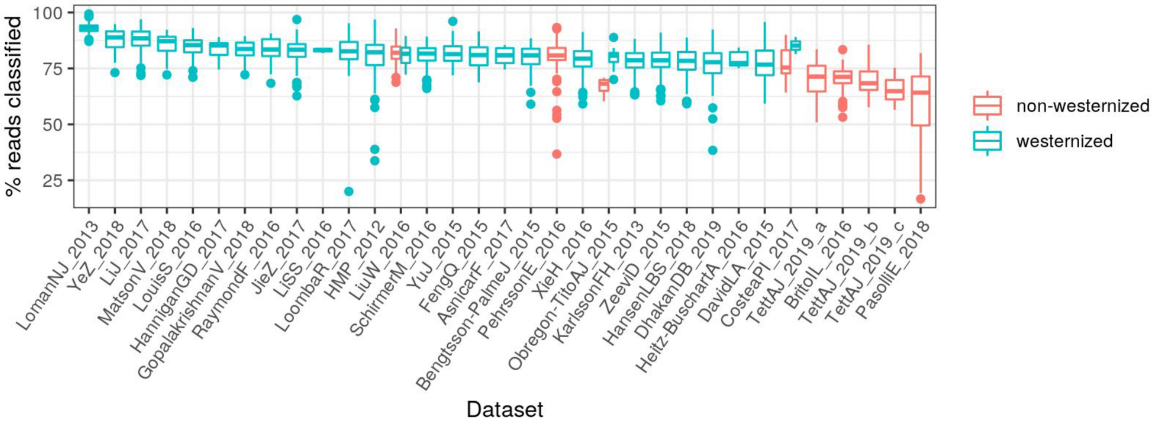


**Figure S1.** The percent of metagenome reads classified by Kraken2 against the GTDB-r89 database, grouped by dataset and colored by westernization status. One million single-end reads per sample were used for the profiling.

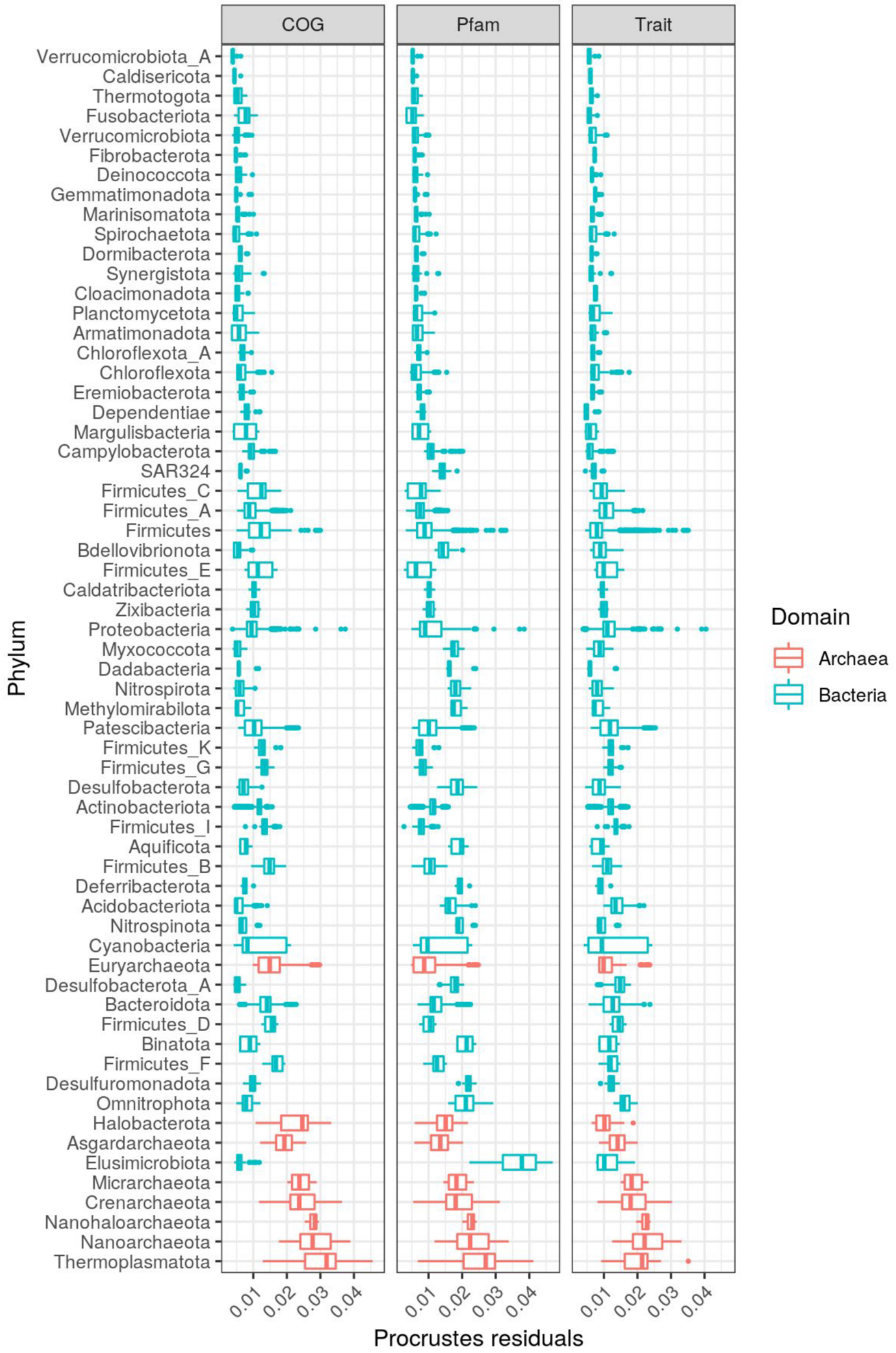


**Figure S2.** Procrustes residuals when comparing the GTDB phylogeny versus trait, COG, or Pfam trees. For clarity, only phyla with ≥10 genomes are shown.

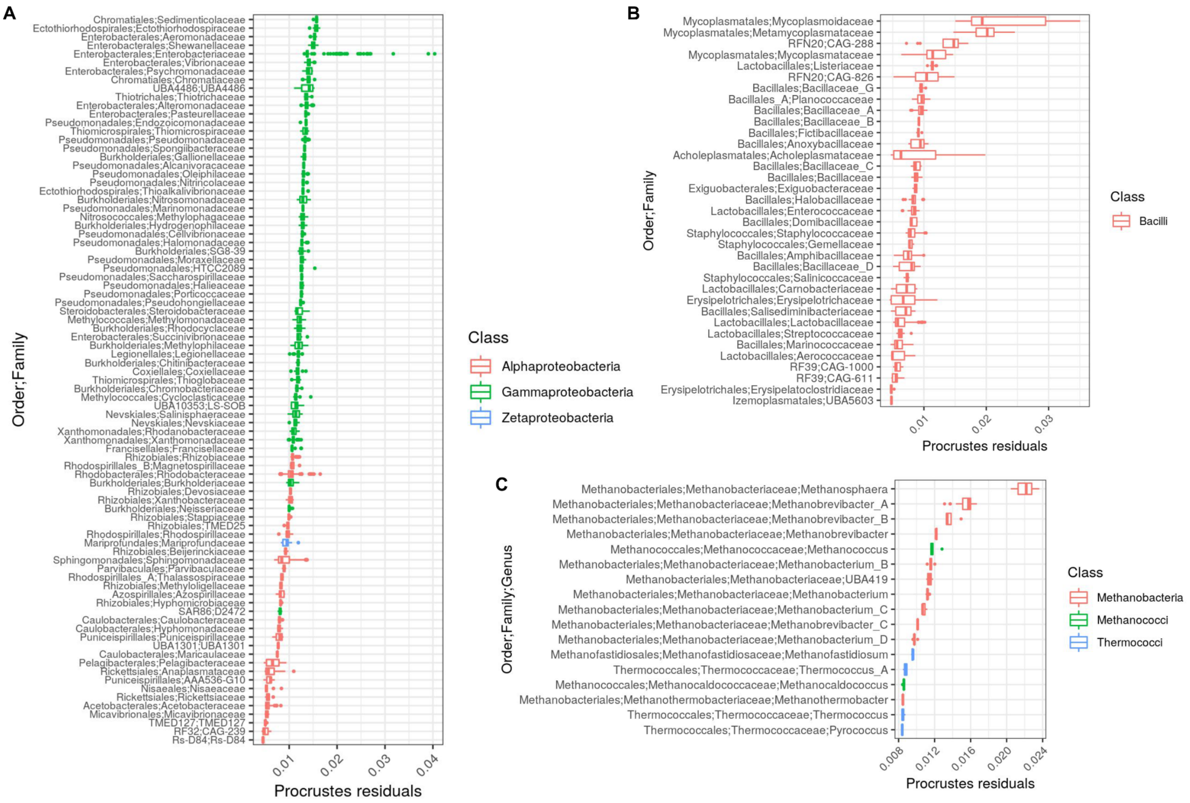


**Figure S3.** *Substantial within-phylum differences in phylogeny-trait congruence.* The boxplots are similar Figure 2 but show Procrustes residuals for families or genera within the phyla: A) *Proteobacteria* B) *Firmicutes* C) *Euryarchaeota*.

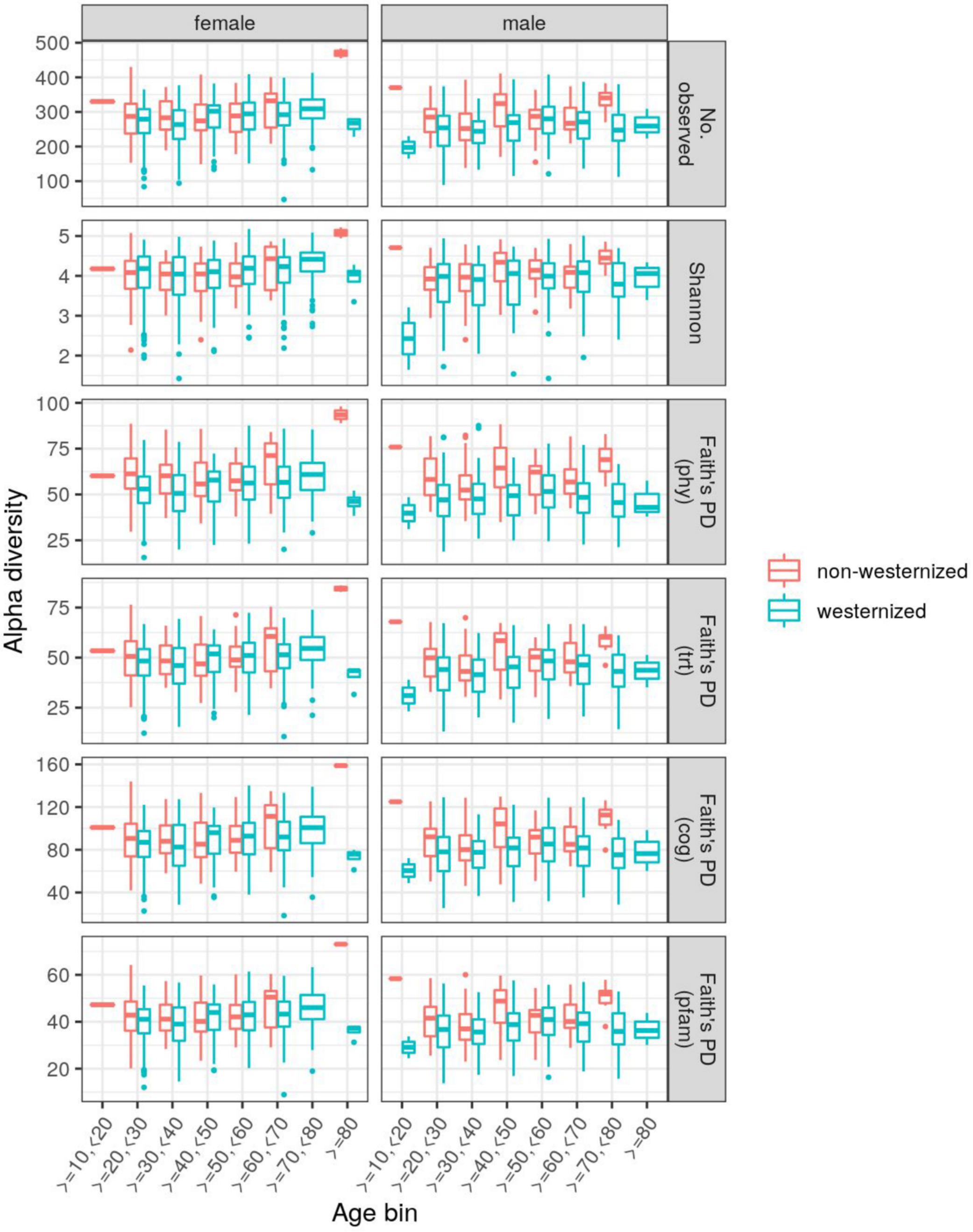


**Figure S4.** Alpha diversity by age, gender, and westernization for all samples with westernization, gender, and age metadata (*n* = 1843). "trt" stands for "traits", while "phy" stands for "genome phylogeny".

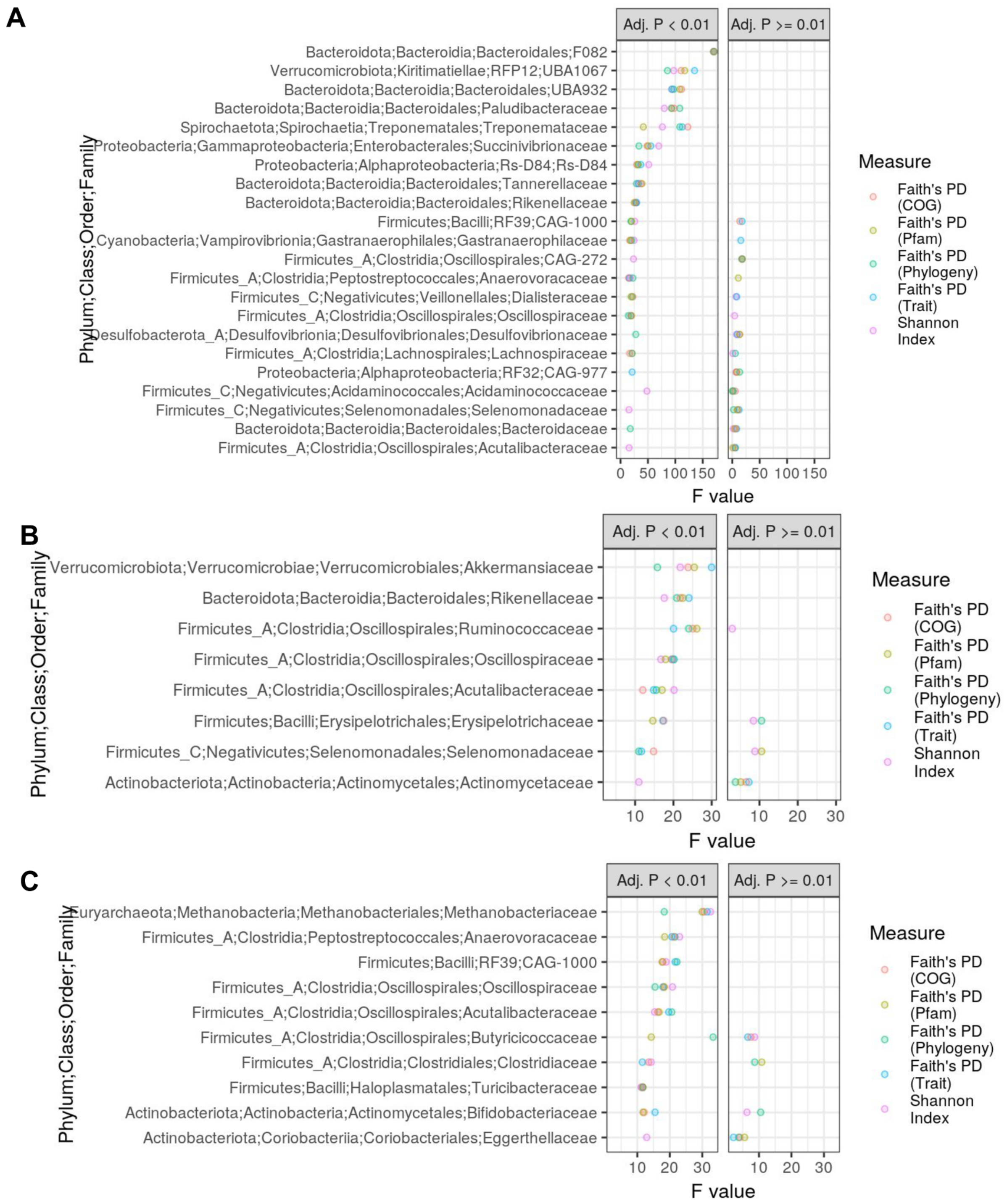


**Figure S5.** *Model effect sizes vary depending on the alpha diversity measure.* Results for linear mixed effects models assessing the association between family-level alpha diversity in relation to A) westernization, B) gender, and C) age (number of samples = 1843; number of studies = 17). Note that "dataset" was included as a random effect. Age was $\log_2$-transformed. Alpha diversity was only calculated for families with >1 species. For clarity, families are shown only if ≥1 mixed model produced an adjusted $P < 0.01$.

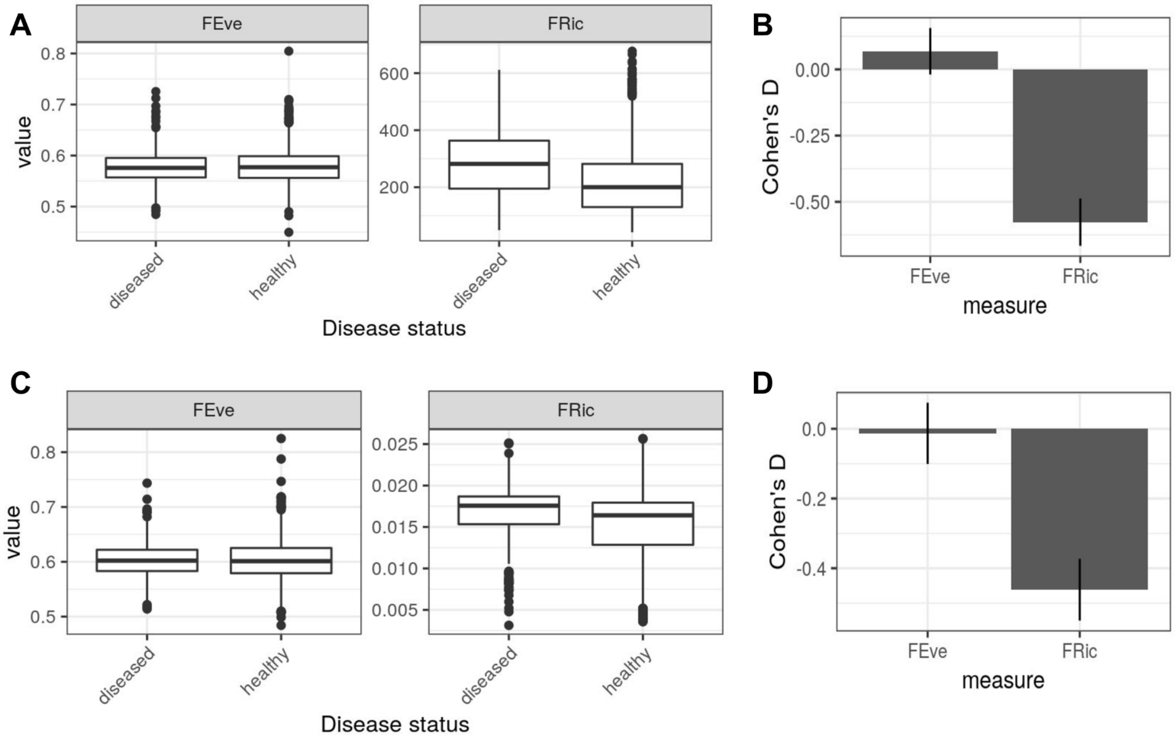


**Figure S6.** *Functional richness but not functional evenness differs by disease status.* The distribution of functional evenness (FEve) and functional richness (FRic) by disease status. Functional diverversity was calculated from either A-B) inferred traits or C-D) COG categories. The bar plots show the effect sizes (Cohen's D) for differences in functional diversity between diseased and healthy samples. The line ranges denote 95% confidence intervals.

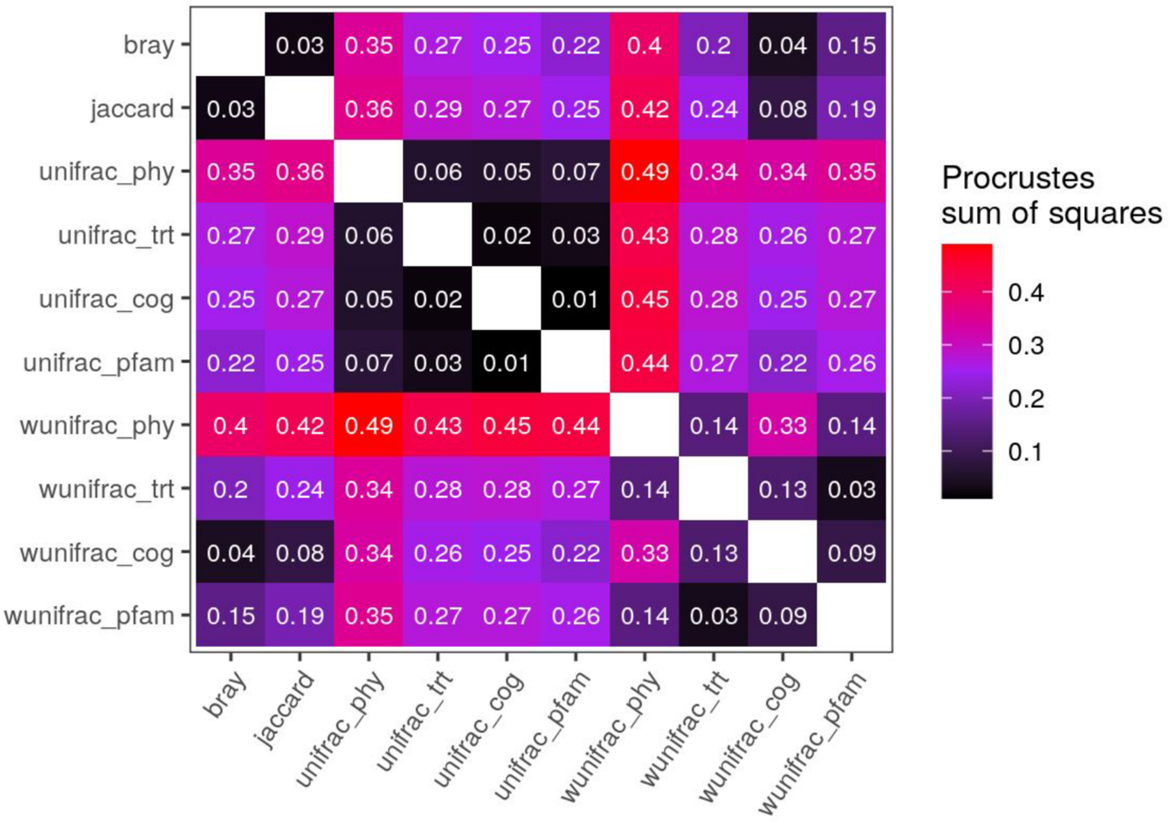


**Figure S7.** *Beta diversity measures markedly differ in how they represent inter-community similarity.* Procrustean comparison of the PCoA ordinations shown in Figure 6. Procrustean sum of squares (higher values indicate greater dissimilarity between ordinations). All pairwise comparisons were significantly different as determined by a protest permutation analysis (999 permutations, adj. $P < 0.05$). Note that the matrix is symmetric along the diagonal. “trt”, “phy”, “unifrac”, and “wunifrac” stand for “traits”, “genome phylogeny”, “unweighted UniFrac”, and “weighted UniFrac”, respectively.

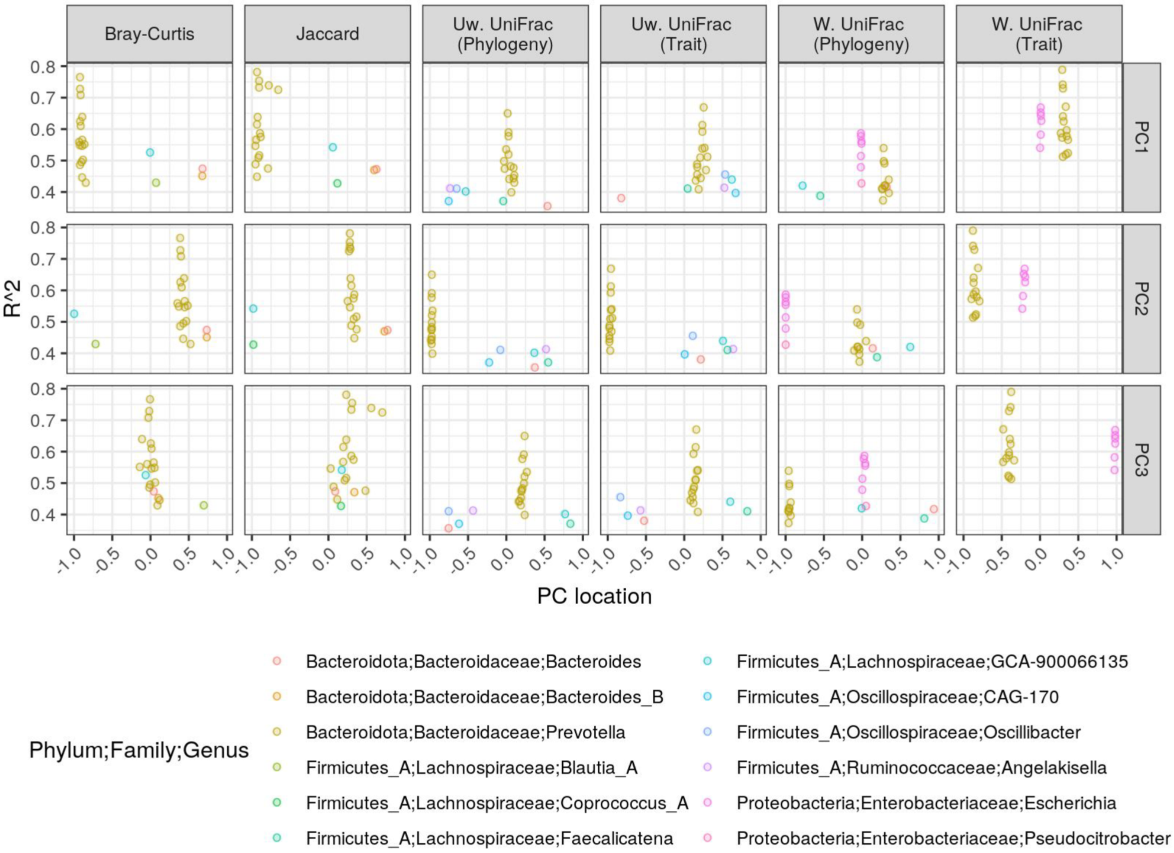


**Figure S8.** The same as Figure 7A, but species (points) are colored by genus level classifications.

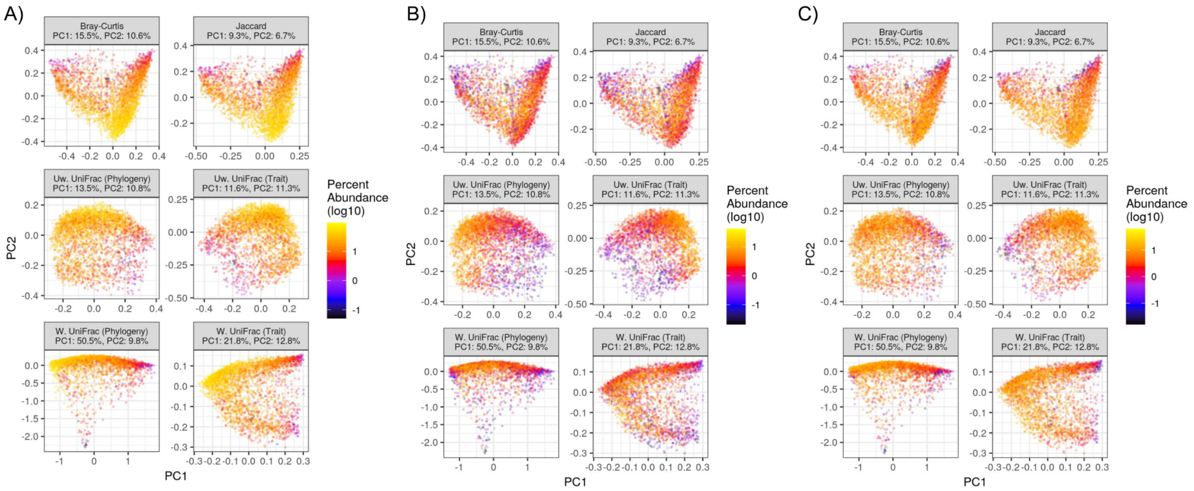


**Figure S9.** The PCoA ordinations as show in Figure 6A, but the points (samples) are colored by the abundance of A) *Lachnospiraceae*, B) *Oscillospiraceae*, or C) *Ruminococcaceae* families (each in the *Firmicutes A* phylum), respectively.

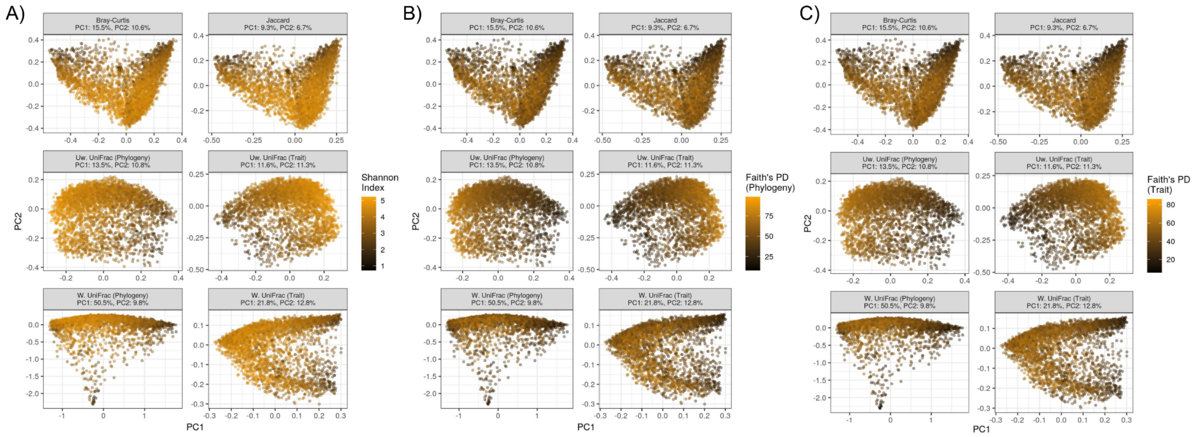


**Figure S10.** PCoA ordinations as shown in Figure 6A, but the samples are colored by the alpha diversity measures: A) Shannon Index, B) Faith’s PD based on the genome phylogeny, and C) Faith’s PD based on the trait similarity dendrogram.
